# Detection of de novo copy number deletions from targeted sequencing of trios

**DOI:** 10.1101/252833

**Authors:** Jack M. Fu, Elizabeth J. Leslie, Alan F. Scott, Jeffrey C. Murray, Mary L. Marazita, Terri H. Beaty, Robert B. Scharpf, Ingo Ruczinski

**Author notes:** To whom correspondence should be addressed: Department of Biostatistics, Johns Hopkins Bloomberg School of Public Health, 615 North Wolfe Street, Baltimore MD, 21205, USA.

## Abstract

*De novo* copy number deletions have been implicated in many diseases, but there is no formal method to date however that identifies *de novo* deletions in parent-offspring trios from capture-based sequencing platforms. We developed Minimum Distance for Targeted Sequencing (MDTS) to fill this void. MDTS has similar sensitivity (recall), but a much lower false positive rate compared to less specific CNV callers, resulting in a much higher positive predictive value (precision). MDTS also exhibited much better scalability, and is available as open source software at github.com/JMF47/MDTS.

## Background

Copy number variants (CNVs) are a major contributor of genome variability in humans [1], and frequently underlie the etiology of disease [2, 3, 4, 5, 6]. *De novo* CNVs, especially *de novo* deletions, are of interest as they have the potential to play a functional role in the genesis of a disease phenotype [7, 8, 9, 10]. Over the last decade, next generation sequencing (NGS) has become routine and widespread [11, 12], permitting the assessment of CNVs based on hundreds of millions of short reads observed in each sample. The advantages of NGS for CNV assessment compared to single nucleotide polymorphism (SNP) arrays include higher and more uniform coverage, better quantitation yielding more precise estimates of DNA copy number, and higher resolution for break point detection [13, 14]. Computational methods to detect CNVs from NGS short reads can generally be categorized into approaches based on discordant read mapping, split read mapping, read depth, *de novo* assembly, or a combination of these approaches [15]. Due to the differences in the attempted capture, methodologies for whole genome sequencing (WGS), whole exome sequencing (WES), and targeted sequencing (TS) platforms differ substantially, with TS and WES platforms primarily relying on read depth [16].

A large number of methods for detecting CNVs in independent samples are available for all types of NGS data [17, 18, 19, 20, 21, 22, 23, 24, 25, 26]. However, there is no method to date that identifies *de novo* CNVs in parent-offspring trios from capture-based TS and WES platforms. For WGS platforms, the software TrioCNV jointly calls CNVs in parent-offspring trios [27] using a hidden Markov model (HMM) with 125 possible underlying states to segment the sequencing data (5 possible underlying states per sample: two-copy deletion, one-copy deletion, normal, one-copy duplication, multiple-copy duplication). Its performance in TS or WES platforms however is not well described. In CANOES, also HMM based, inference for *de novo* copy number deletions in TS and WES data is obtained post-hoc by comparing single-sample derived CNV calls. For each sample of the trio, the observed read counts are modeled using negative binomial distributions, and the respective variances are estimated using a regression-based approach based on selected reference samples [28]. However, such approaches do not fully leverage the Mendelian relationship between parents and offspring to delineate *de novo* CNVs. The loss of statistical power for delineating *de novo* CNVs by post-hoc methods has been demonstrated previously in CNV calls from SNP array data [29, 30].

The motivating example in this manuscript is a targeted re-sequencing study we recently carried out in 1,409 Asian and European case-parent trios ascertained by non-syndromic orofacial cleft probands, targeting 13 regions previously implicated in candidate genes and genome-wide association studies (GWASs) [31]. The study successfully confirmed 48 *de novo* nucleotide mutations, and provided strong evidence for several specific alleles as contributory risk alleles for non-syndromic clefting in humans. Choosing two of these nucleotide *de novo* variants for functional assays, we showed one mutation in PAX7 disrupted the DNA binding of the encoded transcription factor, while the other mutation disrupted the activity of a neural crest enhancer downstream of FGFR2 [31]. However, for the majority of trios, we were not able to identify a genetic cause underlying the proband’s oral cleft. Since *de novo* deletions have previously been shown to underlie oral cleft risk [32, 33, 34], we speculated that in addition to *de novo* nucleotide variants, *de novo* deletions in the 13 targeted regions also contribute to clefting for some of our trio’s probands.

In this manuscript, we present a novel method to delineate *de novo* deletions from TS of trios. We propose a novel capture-based definition of targets (using average read depth as the defining metric for bins underlying the algorithm, instead of using a uniform number of base pairs), normalize copy number counts using the entire study population, and utilize a “minimum distance” statistic based on normalized read count summaries, aiming to further reduce shared sources of technical variation between offspring and parents within a trio. We characterize the sensitivity, specificity, and PPV of MDTS on simulated data to benchmark its performance relative to the closest existing methods TrioCNV [27] and CANOES [28]. We show that properly addressing the capture in TS data is critical, and thus, methods specifically developed for WGS data (e.g., TrioCNV) do not perform well for TS data. Compared to CNV callers designed for capture based sequencing data that do not exploit the family design (e.g., CANOES), MDTS has similar sensitivity but a much lower false positive rate, resulting in a much higher PPV. In the analysis of the 6.7Mb TS oral cleft data, which identified one *de novo* deletion in the gene TRAF3IP3 (a suspected regulator of IRF6), MDTS also exhibited much better scalability.

## Results

### The MDTS algorithm

Our novel MDTS introduces two novel algorithmic aspects. First, MDTS employs bins of varying sizes based on read depth (Figure 1, A–D) as compared to the common standard of using uniform, non-overlapping bins defined by the number of nucleotide base pairs (default in CANOES: 200bp capture based data; default in TrioCNV: tiled, non-overlapping 200bp bins). Second, MDTS fully exploits the trio design to infer *de novo* deletions (Figure 1, E–G), as compared to processing the trio samples separately and carrying out post-hoc inference. To demonstrate that both of these algorithmic features are important in the delineation of *de novo* deletions, and to quantify their relative contributions to sensitivity, specificity, and PPV, we compare the default implementations of MDTS and CANOES in the following section, plus MDTS based on the uniform, non-overlapping 200bp “probe-based” bins (MDTS:p) and CANOES based on the non-uniformly sized “MDTS bins” (CANOES:b).

**Figure 1:**
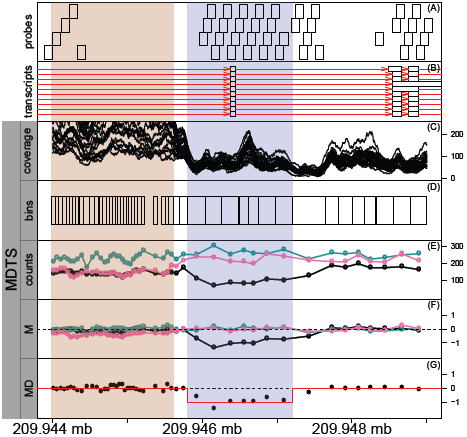
Schematic outline of the MDTS algorithm. Schematic flowchart of the MDTS method, from bin to CNV delineation. **(A)** Design probes in the genomic regaion between 209.944 Mb and 209.948 Mb of chromosome 1. The probes are approximately 120bp long, and often overlap by 60bp **(B)** Transcripts (red lines) from the GencodeV27 annotation. Ten transcripts of TRAF3IP3 contain the exon (white boxes) in the region shaded blue. **(C)** Basepair coverage (read depth) derived from the 25 samples randomly selected to calculate MDTS bins. The region indicated by the rose color was flagged by MDTS for high variability. **(D)** MDTS bins calculated from read depth, leading to wider bins when coverage is low (and vice versa). **(E)** Read depths for the MDTS bins among the three DS10826 family members (proband in black). **(F)** Normalized counts (M-scores) for the three DS10826 family members. **(G)** The minimum distance for family DS10826, and the outcome from CBS segmentation (red line), inferring a candidate *de novo* deletion.

### Simulation Study

MDTS and CANOES produced somewhat similar results for sensitivity (recall) among *de novo* deletions of 1kb or larger, while CANOES had better sensitivity for very small *de novo* deletions. As expected, the algorithms using small uniform and non-overlapping 200bp bins (“probe-based bins”) faired slightly better for small *de novo* deletions, while using read depth based bins (“MDTS bins”) had higher sensitivity for 1kb *de novo* deletions or larger (Figure 2A, Supplementary Table 1). These findings remained the same under other definitions of “overlap” between called and simulated deletions (Supplementary Figure 1). Very pronounced differences were observed with regards to the number of false positive identifications. Depending on size, up to 10% of inherited deletions were incorrectly identified as *de novo* by CANOES using the default 200bp bins (increasing to about 15% for CANOES:b, i.e. when using read depth based bins), while MDTS was extremely robust towards this type of mistake. This was also true when uniform and non-overlapping 200bp bins were used in the MDTS algorithm (e.g., MDTS:p), highlighting the importance of fully exploiting the trio design when inferring *de novo* deletions (Figure 2B, Supplementary Table 2).

**Figure 2:**
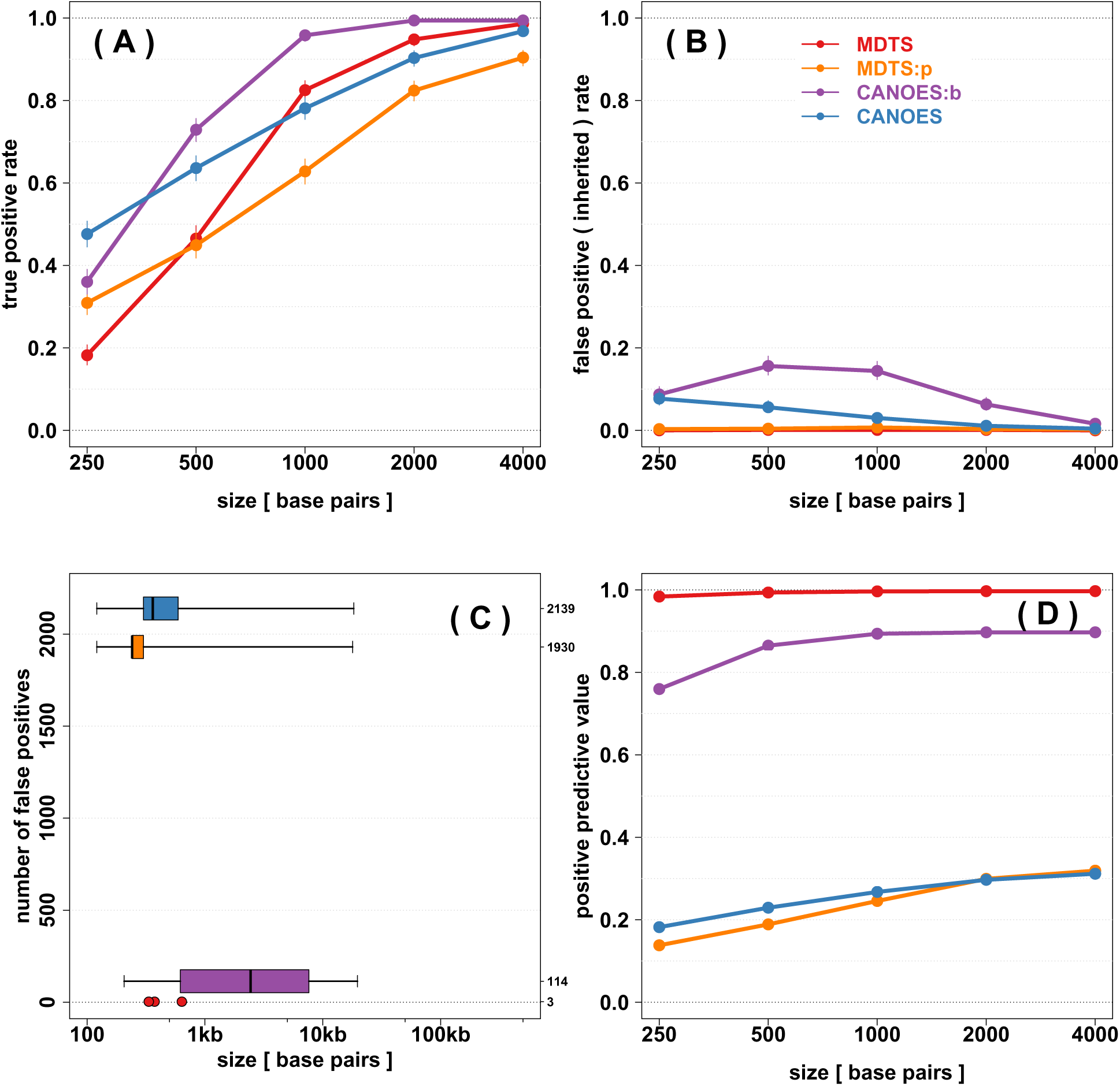
Simulation results to assess sensitivity, specificity, and positive predictive value of four different algorithms to infer de novo deletions. **(A)** True positive rate (sensitivity, y-axis) among 1,000 iterations for simulated *de novo* deletions of various sizes (x-axis). Point estimates are shown as circles together with Binomial 95% confidence intervals. **(B)** False positive rate (specificity) among 1,000 iterations for simulated inherited deletions of various sizes. **(C)** Number of additional false positive identifications from the simulation experiment (y-axis), with length distribution on the logarithmic scale (x-axis) shown as boxplots. MDTS with the newly defined bins only produced three additional false positves, which are shown as points. **(D)** Positive predictive value based on the true positive rate in panel **(A)** and the false positives in panel **(C)**. Colors indicate the algorithms. MDTS and CANOES refer to the respective algorithms as implemented, MDTS:p refers to MDTS based on the uniform, non-overlapping 200bp “probe-based” bins, CANOES:b refers to CANOES based on the non-uniform read depth based bins.

In addition, MDTS incorrectly identified 3 small *de novo* deletions of 334, 374, and 637 base pairs in this simulation study, while CANOES yielded 2,139 false positives with a median width of 361 base pairs (ranging from 121 to 18,339 base pairs). This number was reduced to 114 false positives when instead our proposed read depth based bins were used in the CANOES algorithm (e.g., CANOES:b), but these inferred deletions were generally larger in size with a median width of 2,440 base pairs and a range of 206 to 19,709 base pairs. The importance of using read depth based bins in the algorithms to control false positive identifications was evident, as MDTS built on probe-based coverage (MDTS:p) also faired a lot worse than MDTS (Figure 2C, Supplementary Table 3). These differences in the numbers of false positive identifications observed among these algorithms also resulted in substantial differences when estimating the PPV. The almost complete absence of false positive identifications in MDTS resulted in PPVs approaching 100%, while CANOES did not exceed 33% even for the large *de novo* deletions. CANOES:b on the other hand achieved about 90% PPV, highlighting the importance to use read depth based bins (Figure 2D, Supplementary Table 4).

As expected, TrioCNV did not perform well in the simulation study due to its design for WGS (i.e. non-capture) data. TrioCNV with default 200bp genomic bins was unable to detect any *de novo* deletions, and TrioCNV with MDTS bins only achieved at most 2% sensitivity even for the larger deletions.

### Oral Cleft Case Study

Of the full complement of 4,227 samples, 3,054 samples in 1,018 case-parent trios passed sequencing quality control metrics. Among these families, the MDTS binning procedure generated 25,305 bins, spanning just over 6.3Mb of the targeted 6.7Mb autosomal region. The bins ranged in size from 19bp to 2,956bp, with a median size of 220bp.

MDTS identified three candidate *de novo* deletions (Table 1). The first candidate spanned a 1.6kb segment on chromosome 1 with an average Minimum Distance of −0.90 across 7 bins, and was strongly supported as a *de novo* deletion by the presence of improperly paired reads spanning this segment (Figure 3, left column). The average read depth for the proband in that region was 714, while a read depth of 1,318 was expected for a copy neutral state. The second candidate region spanned a 1.6kb segment on chromosome 8 with an average Minimum Distance of −0.82 across 7 bins. The average read depth for the proband in that region was 740, which compared to an expected read depth of 1,380 for a copy neutral state, suggesting this proband carried a hemizygous deletion. In contrast to the region on chromosome 1 however, no improperly paired reads spanning this segment were observed, rendering this finding somewhat less conclusive. Thus, this region could also represents a false positive identification (Figure 3, right column). Mendelian inconsistencies among trio genotypes can also indicate a *de novo* deletion [35, 36], while heterozygous genotypes in the proband provide strong evidence against *de novo* deletions, however neither result was observed in this short 1.6kb region for family DS12329 (only 1 variant was reported in the vcf files as 0/0, 0/1, 0/1 for the child and the parents, respectively). The third candidate region spanned a 19kb segment on chromosome 8, with an average Minimum Distance of −0.88 across 74 bins. The apparent deletion in the proband of family DS11025 however was not *de novo*, but inherited from a parent with zero copies (Figure 4, left column). This represents a rather uncommon occurrence, as homozygous deletions typically are only observed for copy number polymorphisms (Figure 4, right column), while the 19kb segment on chromosome 8 was only observed for this one family. In total, MDTS detected and flagged two copy number polymorphic regions, a 7.1kb segment on chromosome 1 and a 3.2kb segment on chromosome 8 (Supplementary Table 5).

**Table 1:** MDTS inferred de novo deletions in the oral cleft data. The region on chromosome 1 (top row) is a genuine *de novo* hemizygous deletion of approximately 1,556 base pairs in the proband of family DS10826, inferred using the minimum distance and supported by aberrantly spaced reads (Figure 3, left column). The region of about 1,557 base pairs near 129.6Mb on chromosome 8 (middle row) likely is a false positive identification, inferred based on read depth and the minimum distance, but not supported by aberrantly spaced reads (Figure 3, right column). The region of about 19kb near 130.1Mb on chromosome 8 (bottom row) stems from an unusal Mendelian event in family DS11025 outside a copy number polymorphism (Figure 4). MD: average minimum distance in the respective regions.

|  | start locus | end locus | size | family | MD |
| --- | --- | --- | --- | --- | --- |
| chromosome 1 | 209,945,655 | 209,947,210 | 1,556 | DS10826 | -0.90 |
| chromosome 8 | 129,614,522 | 129,616,078 | 1,557 | DS12329 | -0.82 |
| chromosome 8 | 130,113,612 | 130,132,753 | 19,142 | DS11025 | -0.88 |

**Figure 3:**
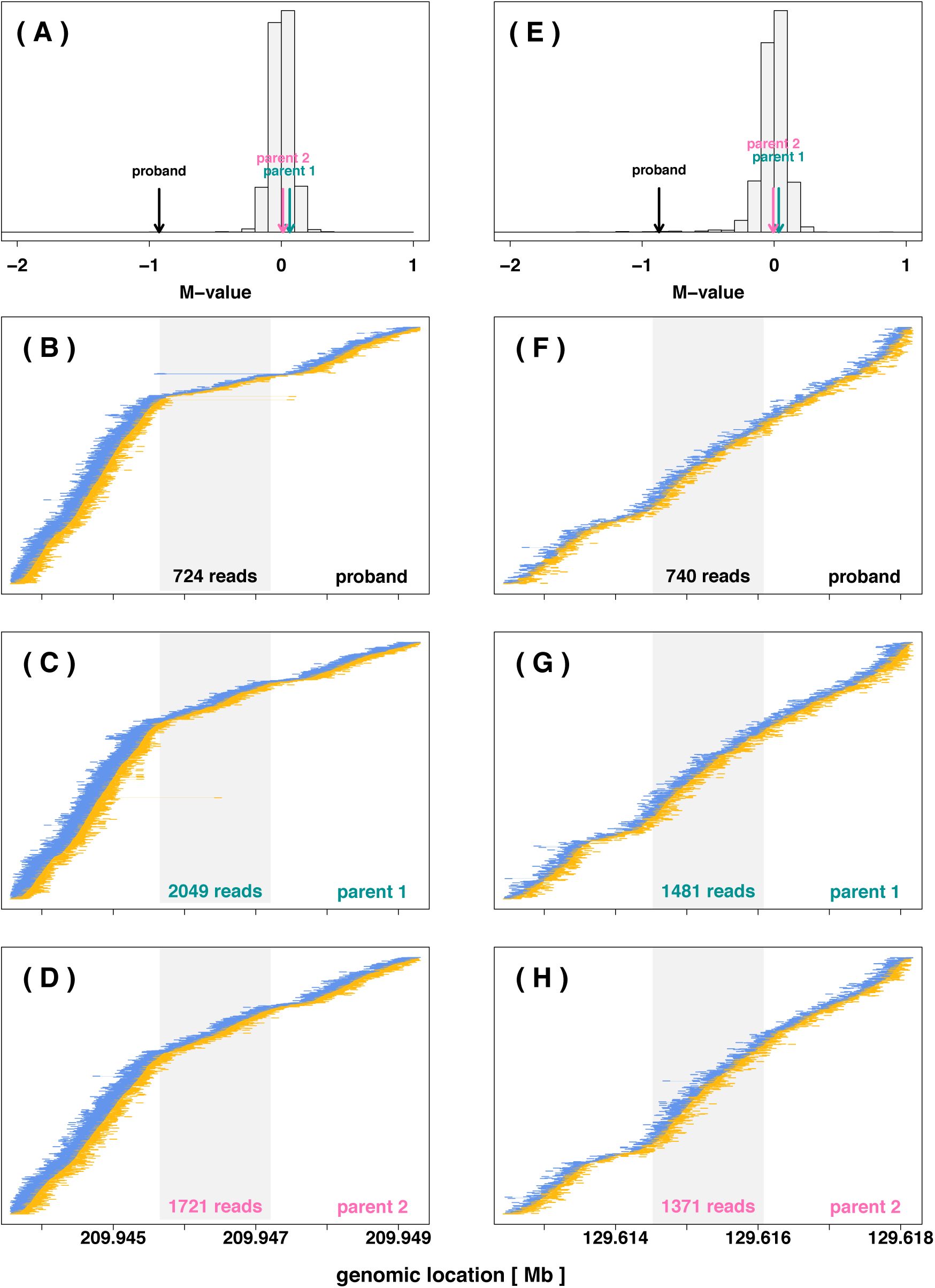
Data underlying inferred de novo hemizygous deletions in two probands. [ **Left Column** ] Evidence for a *de novo* hemizygous deletion on chromosome 1 for the proband in family DS10826. **(A)** The average of the M scores of the proband (-0.93, black arrow), the parents (0.06 and 0.01, green and pink arrows, respectively), and all other subjects (gray histogram) between loci 209,945,655 and 209,947,210 on chromosome 1. The proband’s average of the M scores near −1, compared to the values near zero for all other samples including the parents, is consistent with a *de novo* deletion of one allele in this region. (**B – D**) Read-pairs observed among the members of family DS10826 near the region with the *de novo* hemizygous deletion. The read-pair locations, mapped to the hg19 reference genome, are shown as thick ends connected by thin lines (positive strands shown in yellow, negative strands shown in blue), and sorted by beginning location of mate 1 of the read-pair (e.g. yellow lines are left aligned, blue lines are right aligned). Read-pairs mapped far apart, apparent as a long line, are indicative of a deletion between the ends. A Z-shaped signature of read pairs flanked by such discordant reads as seen for the proband is strong evidence for a 1-copy DNA deletion. The gray region in these panels indicate the inferred 1,556bp hemizygous *de novo* deletion region in the proband’s genome. The number at the bottom of the grey regions in each panel indicates the total number of reads mapped to the inferred *de novo* deletion. [ **Right Column** ] A possible false positive identification of a *de novo* hemizygous deletion on chromosome 8 for the proband in family DS12329. **(E)** The average of the M scores of the proband (-0.87), the parents (0.035 and −0.007, green and pink arrows, respectively), and all other subjects (gray histogram) between loci 129,614,522 and 129,616,078 on chromosome 8. The proband’s average of the M scores near −1, compared to the values near zero for all other samples including the parents, is consistent with a *de novo* deletion of one allele in this region. (**F – H**) Read-pairs observed among the members of family DS12329 near the region with the inferred *de novo* hemizygous deletion. The absence of discordant reads and the Z-shaped signature is evidence against a 1-copy DNA deletion.

**Figure 4:**
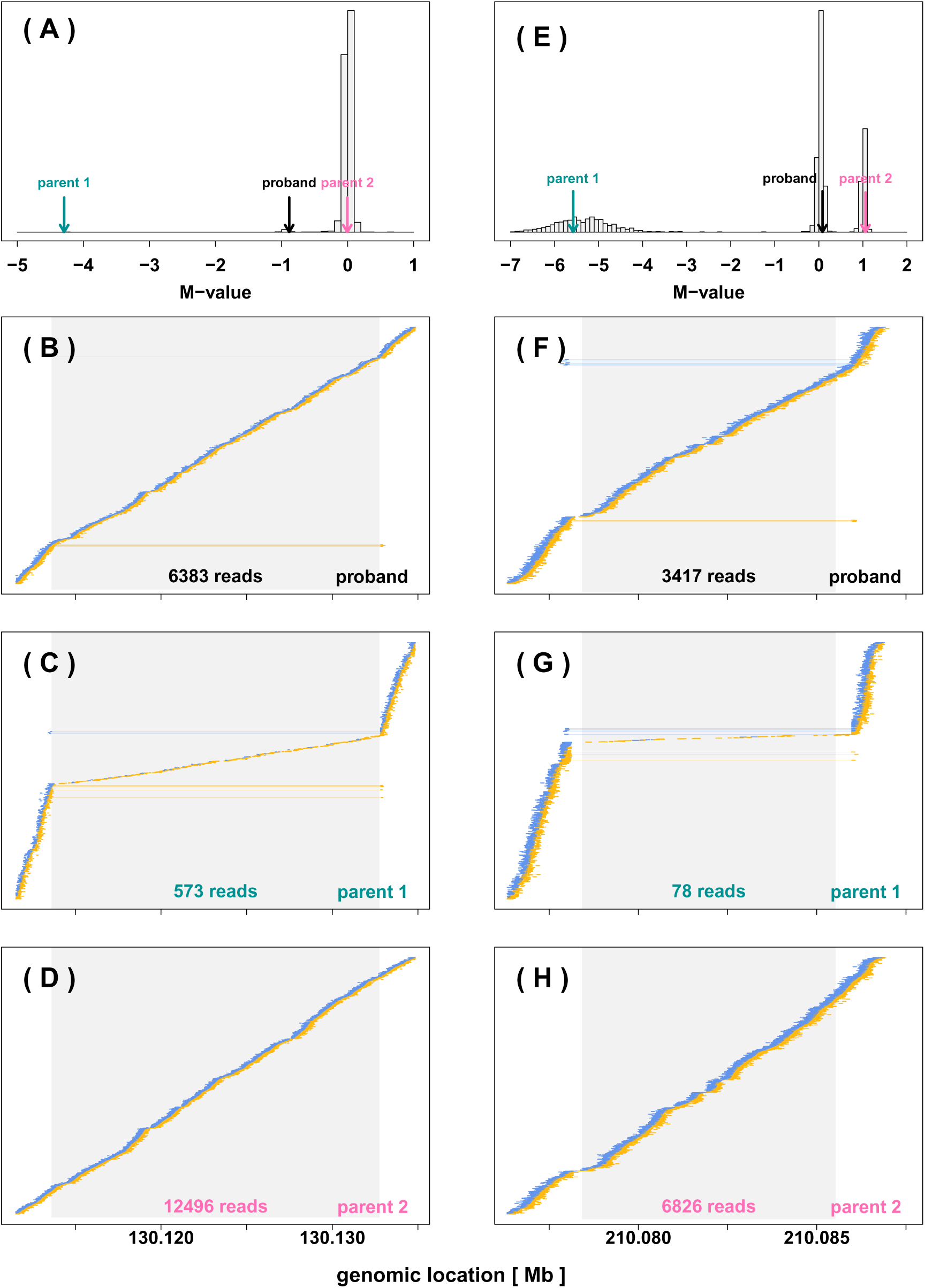
Examples of Mendelian events with a hemizygous deletion in the proband. [ **Left Column** ] A rare Mendelian inheritance event observed on chromosome 8 in family DS11025. (A) The average of the M scores for the proband (-0.88, black arrow) and the parents (-4.3 and −0.01, green and pink arrows, respectively), and all other subjects (gray histogram) between loci 130,113,612 and 130,132,753 on chr8. This is consistent with a hemizygous deletion for the proband, inheriting one copy of the allele from the copy-neutral parent 2, and the deletion from parent 1 showing a homozygous deletion. (B – D) Read-pairs observed among the members of family DS11025 near the region with the inferred Mendelian inheritance event, using the same plotting approach as described in the Figure 3 legend. The Z-shaped signature of a substantial number of read pairs flanked by aberrantly spaced reads seen for the proband again is evidence for a 1-copy (hemizygous) deletion. The Z-shaped signature sandwiching very few (presumably incorrectly mapped) reads for parent 1 is evidence for a 2-copy (homozygous) deletion. The read pairs for parent 2 show a copy-neutral state. The gray region in these panels indicate the inferred 18,956 bp inherited deletion region. The number at the bottom of the grey regions in each panel indicates the total number of reads mapped to the inferred *de novo* deletion. [ Right Column ] A Mendelian inheritance event observed at a copy number polymorphic region on chromosome 1 in family DS11230. (E) The average of the M scores for the proband (0.084, black arrow) and the parents (-5.58 and 1.06, green and pink arrows, respectively), and all other subjects (gray histogram) between loci 210,078,417 and 210,085,527 on chr1. This again is consistent with a hemizygous deletion for the proband, inheriting one copy of the allele from the copy-neutral parent 2, and the deletion from parent 1 showing a homozygous deletion. Due to the polymorphic nature of this region, the initial median normalization failed to correctly center the copy neutral state at zero, which was subsequently inferred by the post-segmentation filter. (F – H) Read-pairs observed among the members of family DS11230 near the region with the inferred Mendelian inheritance event, supporting the inferred 7,111bp hemizygous (homozygous) deletion in the proband (parent 1).

**Figure 5:**
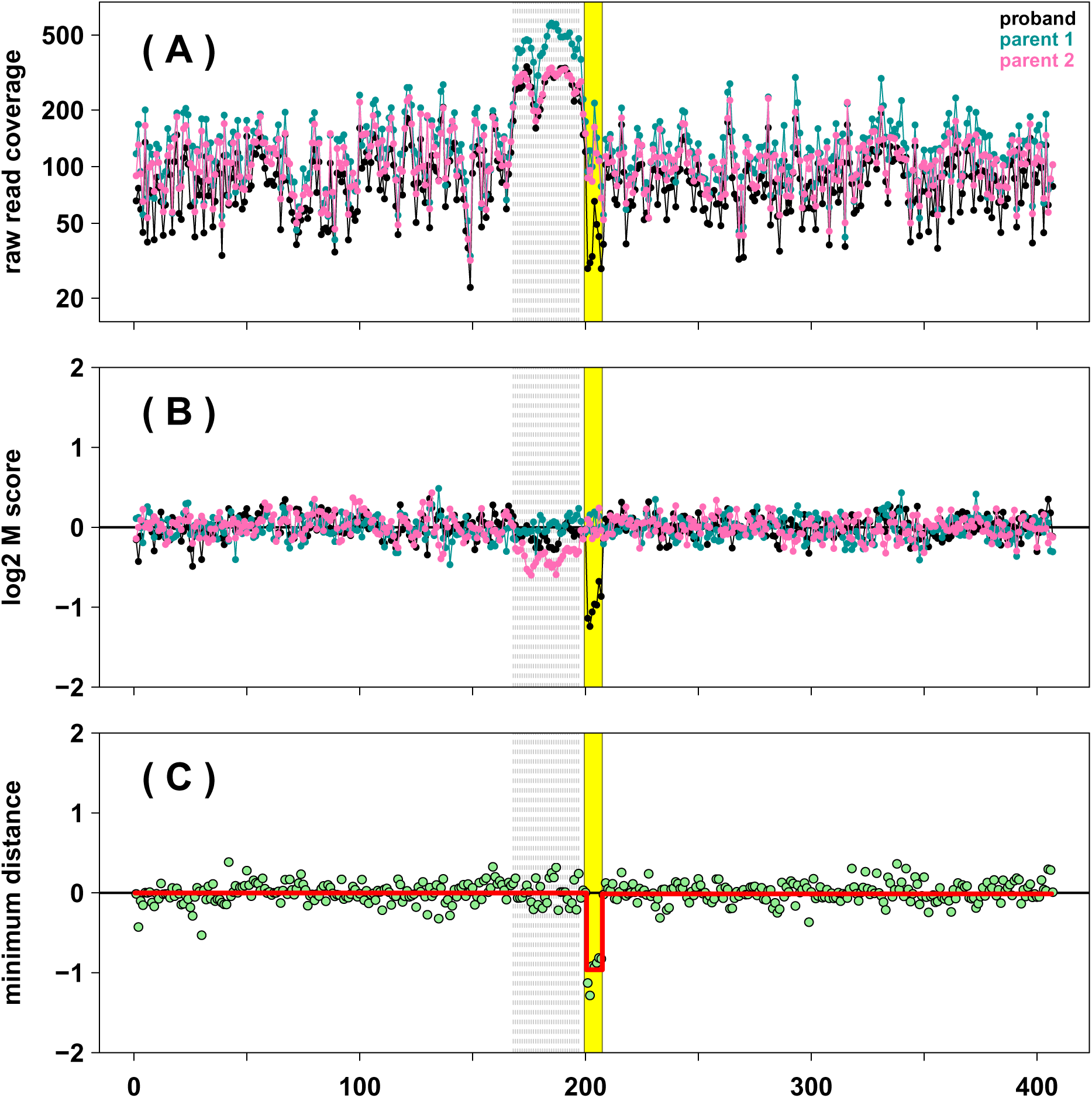
Results obtained using the processing pipeline for a region of 400 bins surrounding the inferred de-novo hemizygous deletion in the proband of family DS10826. **(A)** The coverage of each bin for the proband (black) and the parents parent (green and pink, respectively). **(B)** The M scores for each family member after median centering, GC adjustment, and mappability correction. **(C)** The Minimum Distances for each bin in family DS10826, calculated from the M scores of the proband and the parents. For the region with the inferred de-novo deletion in the proband (yellow segment spanning bins 200–207, corresponding to the genomic loci between 209,945,655 and 209,947,210) the Minimum Distances are close to the expected value of −1, while all other Minimum Distances are centered around zero. The deletion is inferred by Circular Binary Segmentation (solid red line). The gray shaded region spanning 168 through 197 is a region flagged for high variability across M-scores of all samples.

CANOES also identified the true *de novo* deletion in the proband of family DS10826, did not identify the inherited deletion in family DS11025, and did not report the inconclusive MDTS identification in family DS12329. Consistent with the general findings in the simulation study, CANOES also reported a large number of additional *de novo* deletions. In the targeted 6.7Mb region – representing only 0.2% of the genome – the algorithm identified an additional 2,969 *de novo* deletions among the 1,018 families, i.e. about 3 *de novo* deletions per trio on average. Among those 2,969 identifications, 2,702 had a Minimum Distance (calculated from probe-based coverage) outside the [-1.3, −0.7] interval, not consistent with *de novo* deletions (Supplementary Figure 2). The remaining 267 reported *de novo* deletions with Minimum Distances in the interval [-1.3, −0.7] were small (median width 361bp), and none had improper read-pairs spanning the length of the indicated deletion (Supplementary Figure 3). CANOES:b utilizing the MDTS determined bins had the same calls as CANOES reported above for families DS10826, DS11025, and DS12329, but only returned 79 additional *de novo* deletions (though only 28 of those overlapped with any of the 2,969 deletions identified by CANOES). Among those 79 identified deletions, 67 had average Minimum Distances outside the [-1.3, −0.7] interval, and so were inconsistent with *de novo* deletions (Supplementary Figure 4). Among the remaining 12 apparent *de novo* deletions (median width 619bp) with Minimum Distances in the interval [-1.3, −0.7] one is actually an inherited homozygous deletion (Supplementary Figure 5), while the other 11 are located in flagged regions of highly variable normalized depth of coverage (Supplementary Figure 6). TrioCNV with default bins (tiled non-overlapping 200bp bins within the targeted region) did not report any *de novo* deletions among these 1,018 families. In particular, the algorithm failed to identify the true *de novo* deletion on chromosome 1 of the DS10826 proband. TrioCNV with MDTS bins did identify 24 *de novo* deletions, however, 23 of those were actually inherited deletions (Mendelian events) within the chromosome 1 copy number polymorphism (Supplementary Figure 7). The remaining inferred *de novo* deletion supported by only one bin had a Minimum Distance of −0.94, but no improperly mapped read-pairs spanning the deletion which would support a true *de novo* deletion (Supplementary Figure 8). This version of the algorithm also failed to identify the *de novo* deletion on chromosome 1 of the DS10826 proband.

### Scalability

MDTS completed the analysis of the 1,018 oral cleft trios in under 29 hours using a single core, peaking at 15G of memory in the binning step (Supplementary Tables 6 and 7). The run time was cut to less than 6 hours when employing the distributed computing option with 15 cores, albeit at the cost of increasing the peak memory usage to 160G during the counting step. For CANOES, even after editing the supplied R code (which resulted in an almost ten-fold speed-up of the inference), this analysis still required 1,310 hours of CPU times for a single core, but only 14G of memory. TrioCNV, using default parameters except for the distance between adjacent CNVs to be merged and the GC content bin range (see Methods) had a comparable computational footprint to MDTS, requiring 34 CPU hours and 11G of memory to complete the analysis. The usage of MDTS bins slightly reduced the run time for TrioCNV and cut the CANOES CPU time about in half, though the latter was still an order of magnitude slower than MDTS and TrioCNV. MDTS based on probe based bins (MDTS:p) required additional CPU time for the inference compared to the default (MDTS), presumably due to the auto-correlation of the Minimum Distance estimates (resulting from the overlapping design probes) passed to CBS, making break point selection more challenging.

## Discussion

In this manuscript we presented the Minimum Distance for Targeted Sequencing (MDTS) approach for delineating *de novo* copy number deletions simultaneously across multiple trios from TS data. In a simulation study, our approach had a sensitivity competitive with existing methods, but to our knowledge, MDTS is the first caller that rarely generates any false positives. In our simulation study, this approach resulted in a positive predictive value of nearly 100%. We showed this improvement is largely owed to two novel algorithmic features. MDTS employs non-uniformly sized bins based on read depth instead of using uniform, non-overlapping bins defined by the number of nucleotide base pairs, and further, MDTS fully exploits the trio design by using a “minimum distance” statistic to quantify differences in read depths between the offspring and the parents, thereby reducing shared sources of technical variation. We note similar results (equal sensitivity but much improved specificity) were observed for detection of *de novo* deletions based on SNP array data when the Minimum Distance approach was employed, and compared to the results from the trio based PennCNV algorithm [29, 30]. Summarizing the trio data at each locus (probe for SNP arrays or bins for sequencing data) and segmenting these statistics resulted in an estimating procedure with much lower dimensionality than that of a HMM (as used for example in CANOES and TrioCNV). A smaller parameter space is less likely to over-fit, and to generate false positive identifications. Further, fitting a HMM induces an empirical process governing the rate and lengths of these deletions, which may not be realistic as *de novo* deletions are very rare, and could be very small or very large. It should also be noted that MDTS was designed with the sole intent to detect *de novo* deletions in trios, and thus, is much more limited in scope than other CNV callers such as CANOES and TrioCNV (although in principle the MDTS algorithm could also be adapted to detect *de novo* amplifications).

Split reads provide additional compelling evidence for the presence of a copy number deletions, and allow for base pair resolution of break point detection. However, mapping split reads is computationally infeasible for larger deletions unless a candidate has already been identified, and thus, methods based on read depth bins are usually employed to find larger deletions. MDTS is such a method primarily based on read depth, and similar to other read-depth based CNV callers, MDTS has problems identifying very small deletions. In our simulation study, MDTS nonetheless achieved greater than 80% sensitivity for *de novo* deletions of 1kb, and virtually 100% sensitivity for *de novo* deletions of 5kb. We have also implemented functionality allowing for post-hoc inspection of the read ensemble mapped to a region around any putative deletion. In particular the presence of a Z-shaped signature of read pairs flanked by discordant reads - as seen in the suspected IRF6 regulator for the proband of family DS10826 - provides further support for a deletion, and uses information in addition to read depth alone. As the MDTS specificity is very high and *de novo* deletions are rare, the number of candidate deletions to be inspected is low. We queried BAM files to locate split reads that are in the vicinity of a putative deletion. We used SAMtools (samtools.sourceforge.net) to extract split alignments and BLAT (genome.ucsc.edu/cgi-bin/hgBlat) to re-align un-mapped sequences, but were unsuccessful in locating supporting split reads. Thus, no attempts were made to employ LUMPY, arguably the most common CNV caller currently used, to call *de novo* deletions in our data, as its performance heavily relies on such split reads [37]. Further, LUMPY was intended for WGS data and does not account for family structure, thus being less applicable for comparison than TrioCNV and CANOES. Lastly, LUMPY depends on an external read depth caller, which we provide here for TS data in trios.

We also applied our method to 1,305 case-parent trios with 6.7Mb of TS data of regions previously implicated in oral cleft. We detected one *de novo* deletion in the gene TRAF3IP3 on chromosome 1q32 in a Caucasian proband with a cleft lip. TRAF3IP3 is adjacent to IRF6, a gene known to be causal for Van der Woude syndrom (a Mendelian malformation syndrome). Finding only one *de novo* deletion is not too surprising though, as these events are rare, and the MDTS sensitivity is high for deletions larger than 1kb. However, in contrast to single nucleotide variants [38], exact *de novo* mutation rates for copy number variants have not been reported widely. Acuna-Hidalgo et al. [39] estimate one event in 50-100 meiosis for large *de novo* CNVs (in excess of 100kb), but do not give estimates for smaller CNVs citing technical limitations in detecting such events with current short-read sequencing technology. MDTS also returned a second candidate *de novo* region, spanning a 1.6kb segment on chromosome 8. This call was supported by a roughly 50% observed decrease in read depth in this region, in contrast to the region on chromosome 1 however, no improperly paired reads spanning this segment were observed. As no split reads were observed either, an equally confident call whether or not this region harbored a true *de novo* deletion in the proband was not possible. In contrast, one rare inherited deletion identified by MDTS was strongly supported by the observed read depths and improperly paired reads, in addition to two copy number polymorphic regions. It is noteworthy that these two *de novo* deletions as well as the rare inherited deletion identified by MDTS (Table 1) were adjacent to known CNPs on chromosomes 1 and 8, respectively (Supplementary Table 5).

Both CANOES and CANOES:b also identified the true *de novo* deletion in the proband of family DS10826, but did not identify the inherited deletion in family DS11025, and did not report the questionable *de novo* deletion in family DS12329. TrioCNV on the other hand did not perform well due to its design for WGS (i.e. non-capture) data. In our simulation study, CANOES had almost identical sensitivity to MDTS for *de novo* deletions 1kb or larger, which was pushed even higher when using the MDTS bins based on read depth in that algorithm (CANOES:b). In conjunction with a much smaller false positive rate observed (and thus much higher PPV), CANOES:b generally outperformed CANOES in detecting *de novo* deletions (a small caveat however is that CANOES:b was more likely to classify inherited deletions as *de novo*). The reduced number of “hits” from CANOES using our bins compared to the default bins is likely due to our bins avoiding areas where baits were designed, but actual capture was poor. The median MDTS bins size in the oral cleft data analysis was about 160bp, but the size can be controlled by the user. Thus, if detection of smaller *de novo* deletions was a priority, smaller bins could be chosen (which would come at the expense of specificity, naturally).

Scalability of an algorithm is always a concern when working with genomic sequencing data. Even for TS data, CPU demand can be excessive when many samples (or here, many trios) are jointly analyzed. MDTS exhibited much better scalability than CANOES. The oral cleft data analysis was not computationally feasible with the original CANOES code, but we were able to substantially speed up that algorithm by moving a variance-covariance estimation step outside the loop over all trios. Despite running an order of magnitude faster with this tweak, CANOES was still more than an order of magnitude slower than MDTS, and about two orders of magnitude slower than MDTS run multi-threaded. In our opinion it is likely that CANOES was simply not designed with the scale of our oral cleft dataset in mind.

## Conclusions

Our novel method (MDTS) to delineate *de novo* deletions from targeted sequencing of trios fills a specific void in computational approaches for CNV detection. MDTS has similar sensitivity (recall) but a much lower false positive rate compared to related but less specific CNV callers, which results in a much higher positive predictive value (precision). This improvement is owed to using non-uniformly sized bins based on read depth in the algorithm, and fully exploiting the trio design to infer *de novo* deletions. MDTS also has superior scalability.

## Methods

### Samples and Target Region Selection

The original study population included 1,409 case-parent trios comprised of 4,227 individuals of Asian or European ancestry from Europe, the United States, China, and the Philippines (Table S1 in Leslie et al. [31]). Thirteen genomic regions spanning 6.7 Mb were selected for sequencing based on prior association and/or linkage studies, targeting both coding and non-coding sequence at each locus (Table 1 in Leslie et al. [31]).

### Library Preparation, Sequencing, and Alignment

Multiplexed libraries were constructed with 1 mg of native genomic DNA according to standard Illumina protocol with modifications as follows, described in [31]: (1) DNA was fragmented with a Covaris E220 DNA Sonicator (Covaris) to range in size between 100 and 400 bp; (2) Illumina adaptor-ligated library fragments were amplified in four 50 ml PCR reactions for 18 cycles; and (3) solid phase reversible immobilization (SPRI) bead cleanup was used for enzymatic purification throughout the library process, as well as final library size selection targeting 300-500 bp fragments. NimbleGen custom target probes were designed to the target region and hybrid capture on pools of 96 indexed samples per capture was performed. Each capture pool was sequenced on two lanes of Illumina HiSeq for an average of ∼40 Gb per lane or ∼835 Mb per sample. Reads were mapped to the GRCh37-lite reference sequence by BWA v.0.5.912 [40].

### Definition of Bins and M Scores

Due to the prevalence of off-target capture and heterogeneity of coverage within targeted regions, we utilized an empirical approach to define the MDTS bins for computing read depth. Specifically, we randomly sampled 25 subjects and calculated the coverage statistics in each sample across the autosomes. A set of contiguous proto-regions were identified as the set of all basepairs where at least one of the samples had observed coverage of 10x or more. As proto-regions harbored substantial heterogeneity in size depending on both probe density and capture efficiency, the final bins were generated by sequentially partitioned the proto-regions into smaller, non-overlapping regions where the median number of reads across the 25 subsamples was at least 160. Bins were excluded if the average mappability of a bin was less than 0.75, or if the average GC content was outside a “normal” range defined as [0.15, 0.85]. Subsequently, the number of reads overlapping the bins were counted for all samples. The raw count data were organized in a “bin by sample” matrix. We applied a log_2_(count + 1) transformation to reduce skew. Each cell of the matrix was centered by row and column medians. The resulting scores for each sample were further adjusted for average GC content and 100mer mappability of their respective bins, using a locally weighted scatterplot smoother (loess) fit to produce *M* scores, a relative measures of DNA copy number, with an expected value of 0 for a copy-neutral DNA segment, and −1 for a single copy deletion (unless there is a CNP).

### Minimum Distance

To infer *de novo* deletions we utilize the Minimum Distance statistic, previously defined for SNP array data [30]. In brief, at each bin we considered the difference in *M* scores the between the offspring (O) and the father (F), calculated as *M*_*O*_ − *M*_*F*_, and denote this difference as *δ*_*F*_. We calculated the equivalent distance of offspring and mother, and denote this difference as *δ_M_*. The Minimum Distance between parents and offspring at a bin is defined as the smaller of those two differences when comparing their absolute values:

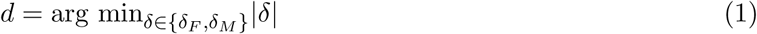

### Filtering and Segmentation

Of the 1,409 families, 383 were removed prior to MDTS bin calculation for experimental design insufficiencies. For these families, the family members were either run in different batches, or did not pass basic quality control as noted by the reporting lab. An additional 8 families were excluded from the analysis based on Minimum Distances summary statistics (lag10 autocorrelation > 0.4 and/or variance > 0.05). Circular Binary Segmentation (CBS) [41, 42], implemented in the Bioconductor package `DNAcopy`, was used for each targeted region to segment the Minimum Distances across the bins for each trio. `CBS` computes a permutation reference distribution of the input Minimum Distances to infer change points for copy number. As this is a random process by default, we fixed a seed `set.seed(137)` in R to ensure reproducibility of our results. We required the minimum number of bins in any segment to be at least 3. In general, default input parameters were used, except using *α* `= 0.001` as the minimum significance required in the CBS t-tests to infer a change point. Further, we allowed change points to be undone when the difference in means was less than 4 standard deviations (`undo.splits=’sdundo’` in conjunction with `undo.SD=4`). Candidate *de novo* hemizygous deletions were identified as regions where the segmented Minimum Distance was within 0.3 of the theoretical value of −1. To reduce the likelihood of false positives based on failures in the normalization process (caused by the existence of CNPs or technical anomalies), regions of high variability were identified as bins where more than 5% of samples had *M* scores outside the interval [-0.5, 0.5]. MDTS reported *de novo* deletions only when more than half of the bins in the candidate region were not flagged.

### Alternative Approach CANOES

This algorithm was designed for capture-based WES and TS data, but the statistical inference does not explicitly take the familial relationship into account. Assessment of *de novo* copy number events in CANOES is based on a post-hoc comparison of the inferred copy number states of the individual samples. The default binning scheme in the algorithm utilizes the bait design coordinates, but MDTS bins can also be used as input. A simple modification had to be made to the CANOES R code, publicly available at www.columbia.edu/∼ys2411/canoes/, to make it scalable for our simulation study and oral cleft data analysis. For large sample sizes (here, n=3,054 in the oral cleft study) the calculation of the n*×*n covariance matrix between bin read counts of samples to locate reference samples for a given individual is computationally very intensive. In the original R code this is carried out for each sample (within the for() loop), but actually has to be carried out only once (outside the for() loop).

### Alternative Approach TrioCNV

In comparison to CANOES, this algorithm explicitly models the proband-parent trio relationship, however was designed for WGS data (i.e., non-capture based sequencing data). The default binning scheme for the inference is based on subdividing the genome into non-overlapping 200bp windows. We restricted these bins to those in the 6.7Mb targeted for sequencing [31]. In the simulation study and the oral cleft data analysis we used the TrioCNV default parameters, with two exceptions: We reduced the value for the argument `min_distance`, which specifies the distance between adjacently called CNVs to be merged, from the default 10,000 to 1,000. We also changed the value for the argument `gc_bin_size`, from its default value of 1 to 2. This value determines the grouping of bins for the estimation of the emission probabilities in the Hidden Markov Model. The default value of 1 did not produce a sufficient number of bins for certain GC values in the capture based data, resulting in JAVA runtime errors thrown.

### Simulation Study

We sampled with replacement 1,000 case-parent trios from the 1,018 families that passed QC. For each instance, we simulated read data based on the TS data for that trio. We first sampled 10 non-overlapping regions among MDTS regions that passed the normalization criterion described above. Of the 10 regions, 5 were designated to harbor *de novo* deletions, and 5 were designated to harbor inherited deletions of sizes 250bp, 500bp, 1,000bp, 2,000bp, and 4000bp. The 5 *de novo* deletion spike-ins were achieved by randomly and independently dropping reads that overlapped the selected regions with probability 0.5 in the probands BAM file. The 5 inherited deletions were generated by randomly and independently dropping reads overlapping the respective regions with probability 0.5 in the BAM files of the proband and one parent. Split reads were not simulated as all methods compared here are based on read-depth. We compared the performances of MDTS, CANOES, and TrioCNV, using default and alternative binning schemes. Specifically, we assessed the performances of MDTS with default read-depth based bins (MDTS), MDTS with probe based bins based on bait design coordinates as defined in CANOES (MDTS:p), CANOES with MDTS bins (CANOES:b), CANOES with default bins (CANOES), TrioCNV with MDTS bins (TrioCNV:b), and TrioCNV with restricted genomic bins as described above (TrioCNV). For CANOES and CANOES:p, the CNV calling was carried out for each family member. Inferred deletions in the proband found to be at least 25% covered by a called deletion in at least one of the parents were deemed to be inherited, otherwise deletions in the proband were considered *de novo*. The spiked-in *de novo* and inherited deletions were considered called if 25% of the deletion was covered by candidates reported. Alternative thresholds of > 0% (any overlap) and 50% (at least half of the deletion was identified) were also considered.

## Declarations

### Ethics approval and consent to participate

The oral cleft study population included case-parent trios recruited by several research groups with samples coming from individuals of Asian or European ancestry from Europe, the United States, China, and the Philippines. Approval for all research work was obtained from the Institutional Review Boards of participating institutions (both US and foreign) and informed consent was obtained from parents of minor children and from all affected individuals old enough to give their own consent. See Leslie et al [31] for details.

### Consent for publication

Not applicable.

### Availability of data and material

MDTS has been submitted for review to the Bioconductor consortium (www.bioconductor.org) and is available as open source software written in the statistical environment R at github.com/JMF47/MDTS. The targeted sequences used in the oral cleft data analysis are available from dbGaP www.ncbi.nlm.nih.gov/gap under accession number phs000625.v1.p1.

## Authors’ contributions

JF, RBS, and IR conceived, designed and implemented the computational methods, and wrote the manuscript. EJL, AFS, JCM, MLM, and THB conceived and carried out the oral cleft targeted sequencing study. All authors read and approved the final manuscript.

## Acknowledgements

We thank the members of the families who participated in the oral cleft study, and the field and laboratory staff who made these methods developments and analyses possible.

## SUPPLEMENTARY MATERIAL

**Supplementary Table 1:** Number of true positive identifications of *de novo* deletions (sensitivity) among 1,000 iterations in the simulation study. MDTS and CANOES refer to the respective algorithms as implemented, MDTS:p refers to MDTS based on the uniform, non-overlapping 200bp “probe-based” bins, CANOES:b refers to CANOES based on the non-uniform read depth based bins.

| size (bp) ⟶ | 250 | 500 | 1,000 | 2,000 | 4,000 |
| --- | --- | --- | --- | --- | --- |
| MDTS | 182 | 465 | 825 | 948 | 986 |
| MDTS:p | 309 | 449 | 628 | 824 | 904 |
| CANOES:b | 360 | 729 | 958 | 994 | 994 |
| CANOES | 476 | 636 | 781 | 903 | 968 |

**Supplementary Table 2:** Number of false positive identifications among the inherited deletions, among 1,000 iterations in the simulation study, for four different algorithms.

| size (bp) ⟶ | 250 | 500 | 1,000 | 2,000 | 4,000 |
| --- | --- | --- | --- | --- | --- |
| MDTS | 0 | 1 | 1 | 1 | 0 |
| MDTS:p | 3 | 4 | 7 | 3 | 2 |
| CANOES:b | 87 | 156 | 144 | 63 | 16 |
| CANOES | 77 | 56 | 30 | 11 | 4 |

**Supplementary Table 3:** The total number N of incorrectly called *de novo* deletions among 1,000 iterations in the simulation study, and median, minimum, and maximum sizes (in nucleotide bases) among those false positives, for four different algorithms.

|  | N | Median | Minimum | Maximum |
| --- | --- | --- | --- | --- |
| MDTS | 3 | 374 | 334 | 637 |
| MDTS:p | 1,930 | 241 | 121 | 17,917 |
| CANOES:b | 114 | 2,440 | 206 | 19,709 |
| CANOES | 2,139 | 361 | 121 | 18,339 |

**Supplementary Table 4:**
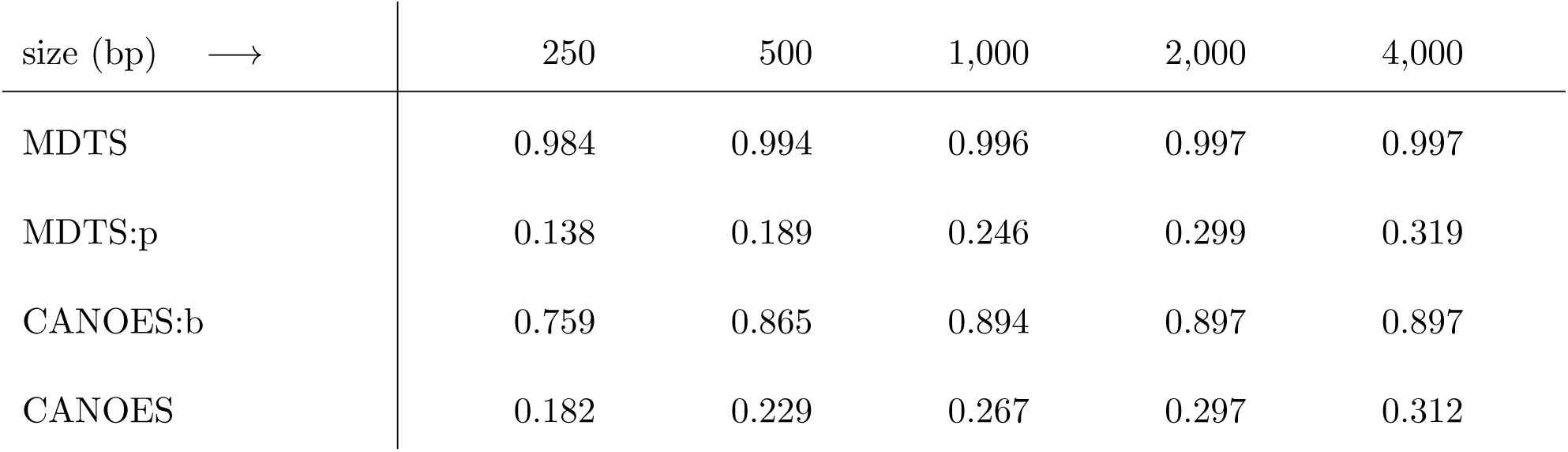
Estimated positive predictive values for the four algorithms.

| size (bp) ⟶ | 250 | 500 | 1,000 | 2,000 | 4,000 |
| --- | --- | --- | --- | --- | --- |
| MDTS | 0.984 | 0.994 | 0.996 | 0.997 | 0.997 |
| MDTS:p | 0.138 | 0.189 | 0.246 | 0.299 | 0.319 |
| CANOES:b | 0.759 | 0.865 | 0.894 | 0.897 | 0.897 |
| CANOES | 0.182 | 0.229 | 0.267 | 0.297 | 0.312 |

**Supplementary Table 5:** Two regions, close to the inferred *de novo* deletions, were highly polymorphic for CNVs in the oral cleft TS data. The data indicate approximate genomic coordinates and the size (in base pairs) of the CNPs, as well as the number of probands with an inherited homozygous (n_hom_) or hemizygous (n_hem_) deletion.

|  | start | end | size | n<sub>hom</sub> | n<sub>hem</sub> |
| --- | --- | --- | --- | --- | --- |
| chromosome 1 | 210,078,417 | 210,085,527 | 7,111 | 297 | 507 |
| chromosome 8 | 129,762,791 | 129,766,015 | 3,225 | 296 | 450 |

**Supplementary Table 6:**
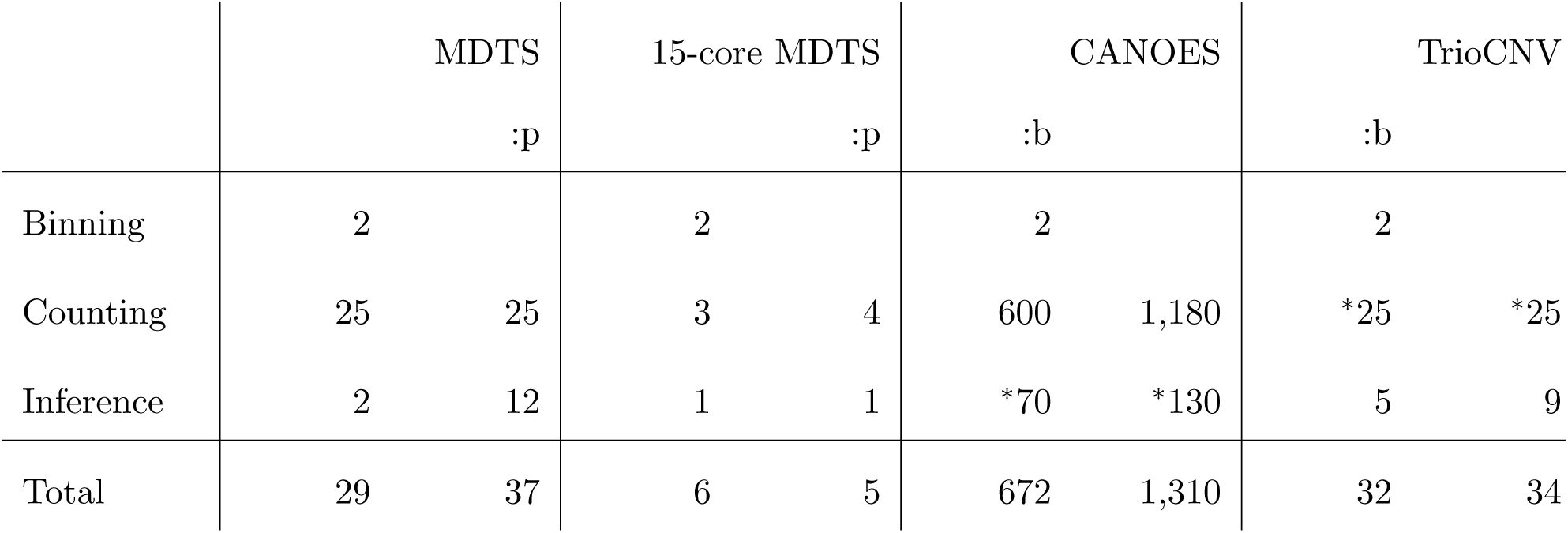
Runtimes (CPU hours, rounded) of MDTS, CANOES, and TrioCNV, and respective modified versions thereof, on the full dataset of 1,018 oral cleft trios, plus runtime for MDTS using an embarrassingly parallel multi-threaded version using 15 cores. Binning refers to the read depth based delineation of MDTS bins using a randomly selected subset of samples, as described in the Methods section. Counting refers to the calculation of read depths for the bins used in the respective algorithms. The asterisk (*) indicates that modifications were made to the publicly available code, described in detail in the Methods section.

**Supplementary Table 7:**
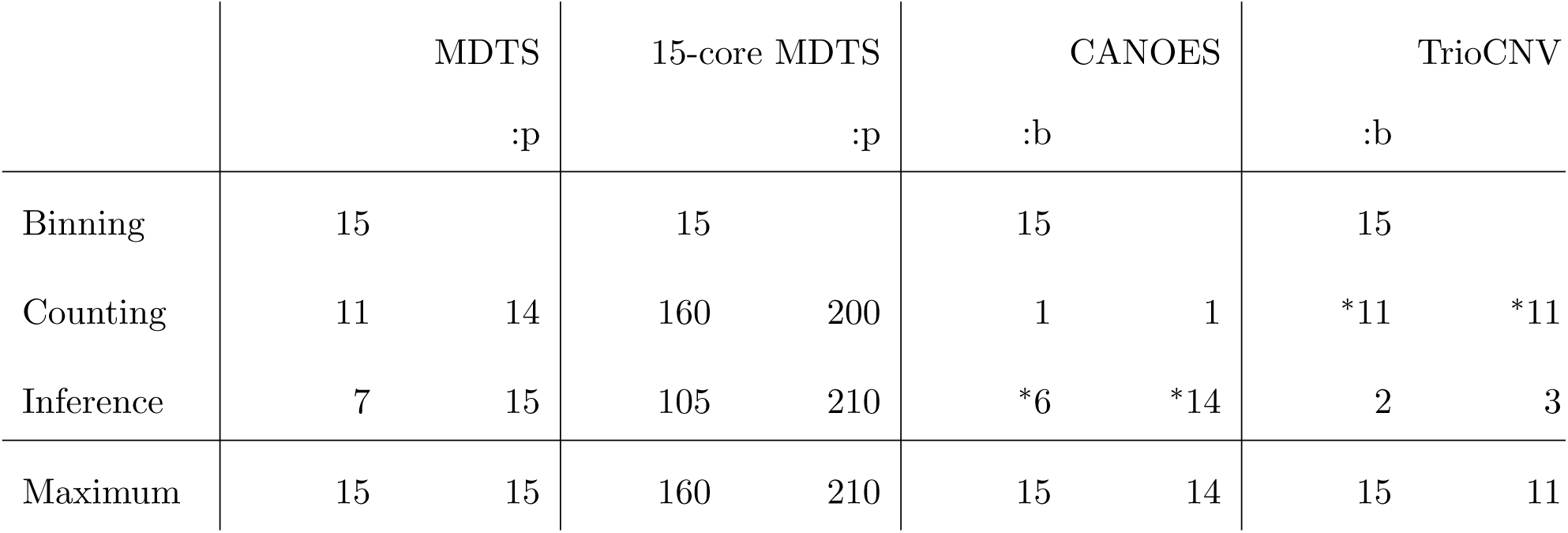
Memory requirements (GB) of MDTS, CANOES, and TrioCNV, and respective modified versions thereof, on the full dataset of 1,018 oral cleft trios, plus memory requirements for MDTS using an embarrassingly parallel multi-threaded version using 15 cores. Binning refers to the read depth based delineation of MDTS bins using a randomly selected subset of samples, as described in the Methods section. Counting refers to the calculation of read depths for the bins used in the respective algorithms. The asterisk (*) indicates that modifications were made to the publicly available code, described in detail in the Methods section.

**Supplementary Figure 1:**
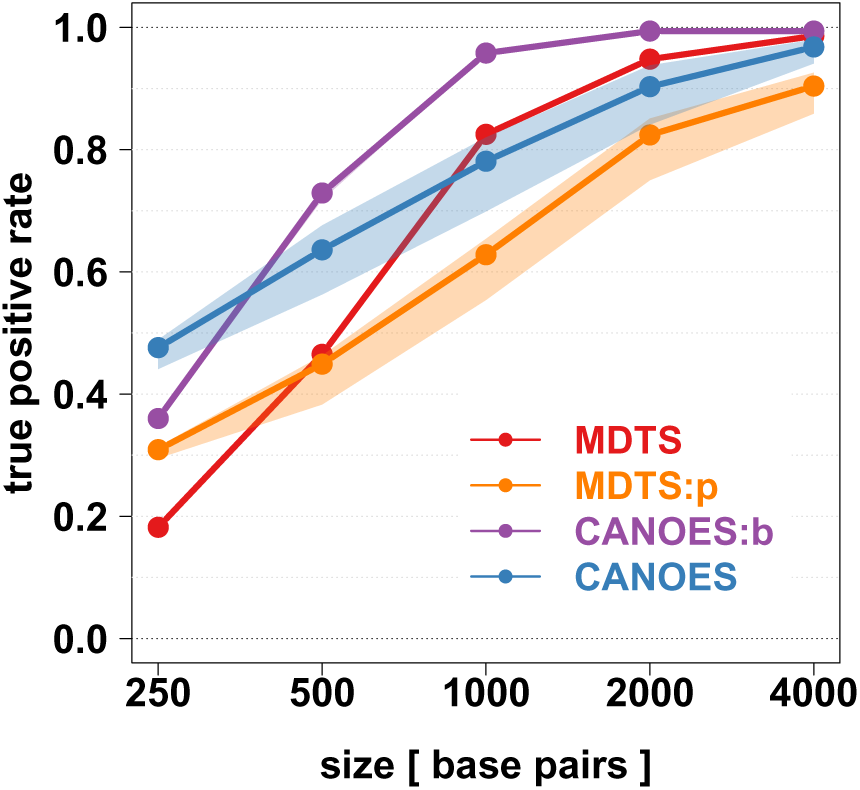
True positive rate (sensitivity, y-axis) among 1,000 iterations for simulated *de novo* deletions of various sizes (x-axis) using different definitions of “overlap” to define true positives, for four different algorithms. The lines show true positive rates using the 25% threshold described in the Methods and shown in Figure 2. The top of the bands result from using a >0% threshold (e.g., any overlap), the bottom of the bands result from a 50% threshold (e.g.m at least half of the deletion was identified).

**Supplementary Figure 2:**
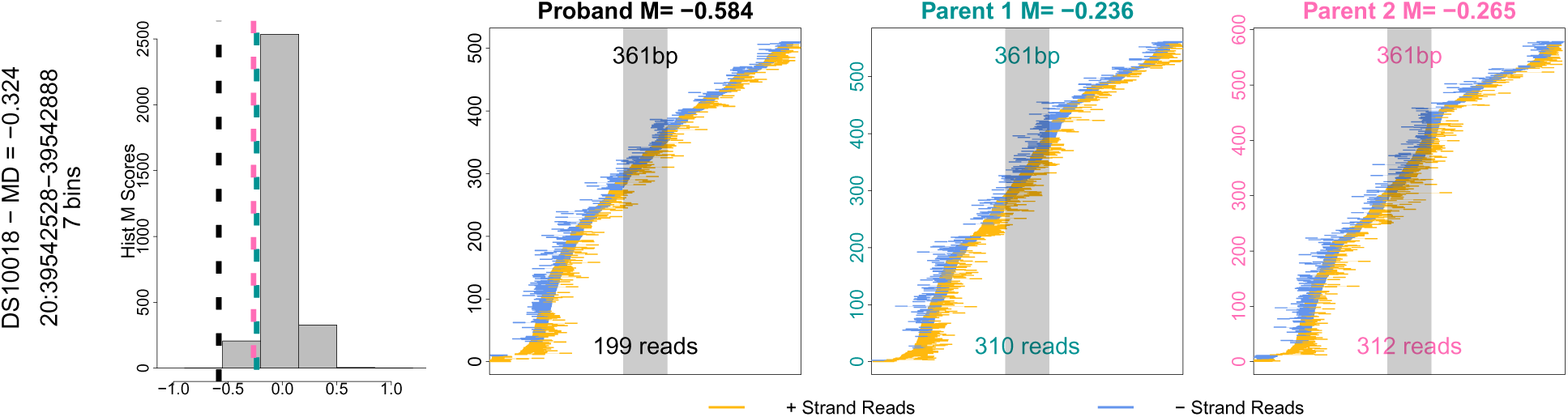
2,702 of the 2,970 trios with CANOES inferred proband *de novo* deletion did not have Minimum Distances consistent with such events. In this example the Minimum Distance was −0.32. The proband does not have discordant read pairs flanking the identified 361bp region.

**Supplementary Figure 3:**
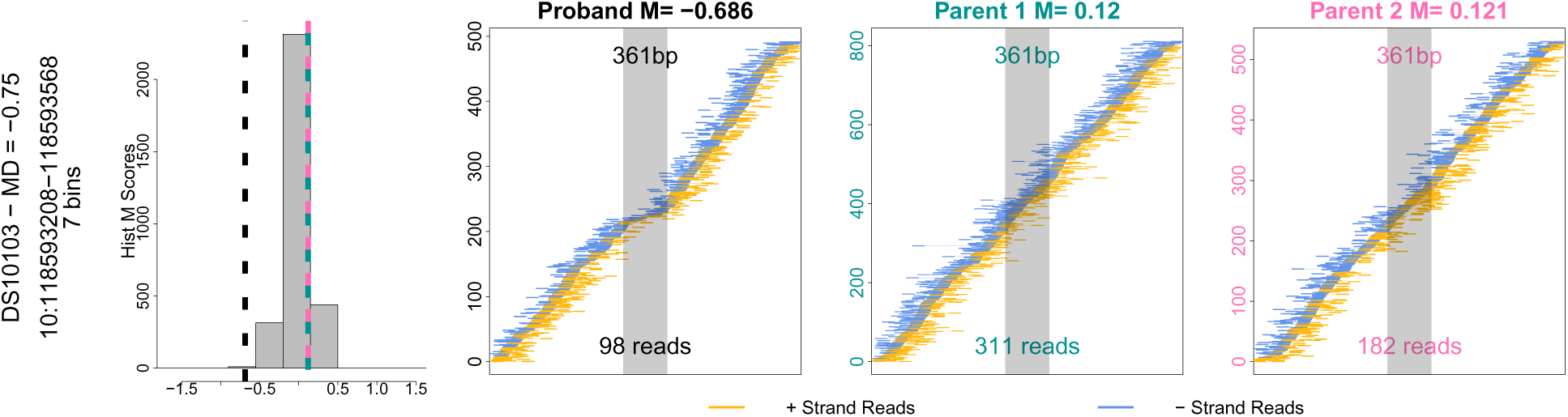
267 trios with CANOES inferred proband *de novo* deletion did have Minimum Distances consistent with such events. In this example the Minimum Distance was −0.75. However, in none of these trios discordant read pairs flanking the identified regions were present.

**Supplementary Figure 4:**
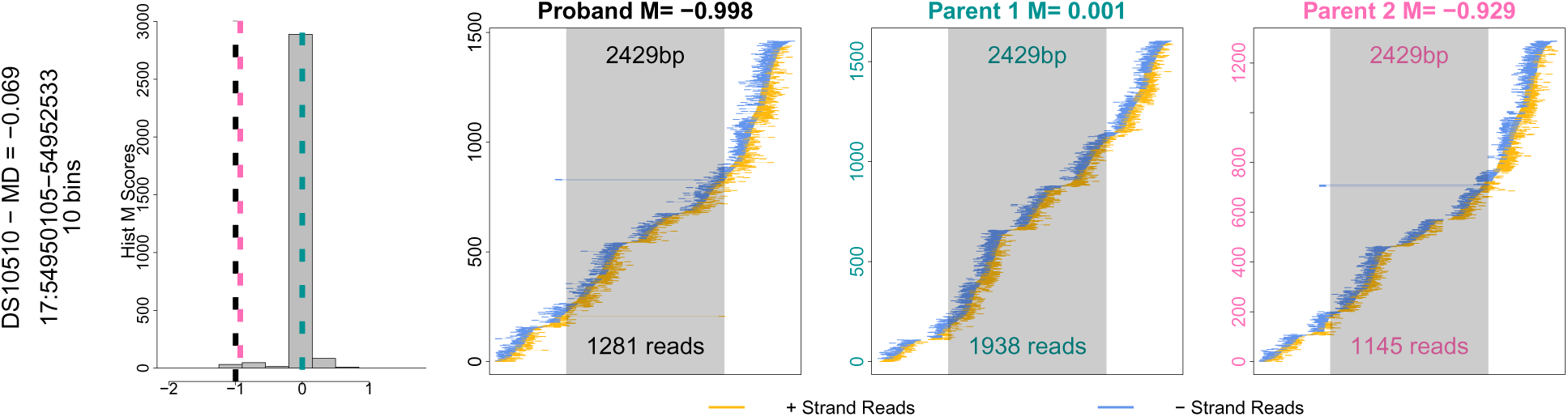
A Medelian event incorrectly called *de novo* by CANOES:b. The M scores (-1.00, 0.00, and −0.93 for the proband and the parents, respectively) and the family Minimum Distance of −0.07 indicate an inherited hemizygous deletion from parent 2. This is further corroborated by the read “Z signatures” in the proband and parent 2.

**Supplementary Figure 5:**
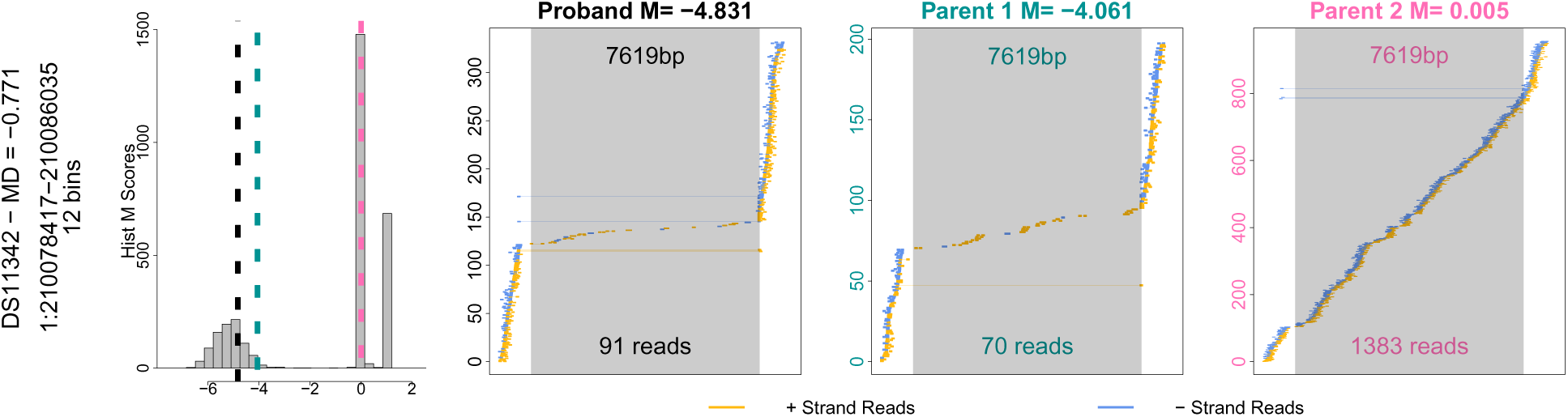
A Medelian event incorrectly called *de novo* by CANOES:b. The M scores (-4.83, −4.06, and 0.01 for the proband and the parents, respectively) and the family Minimum Distance of −0.77 indicate an inherited homozygous deletion in the proband, from one homozygous parent (1) and one hemizygous parent (2). This is further corroborated by the read “Z signatures” in the individuals.

**Supplementary Figure 6:**
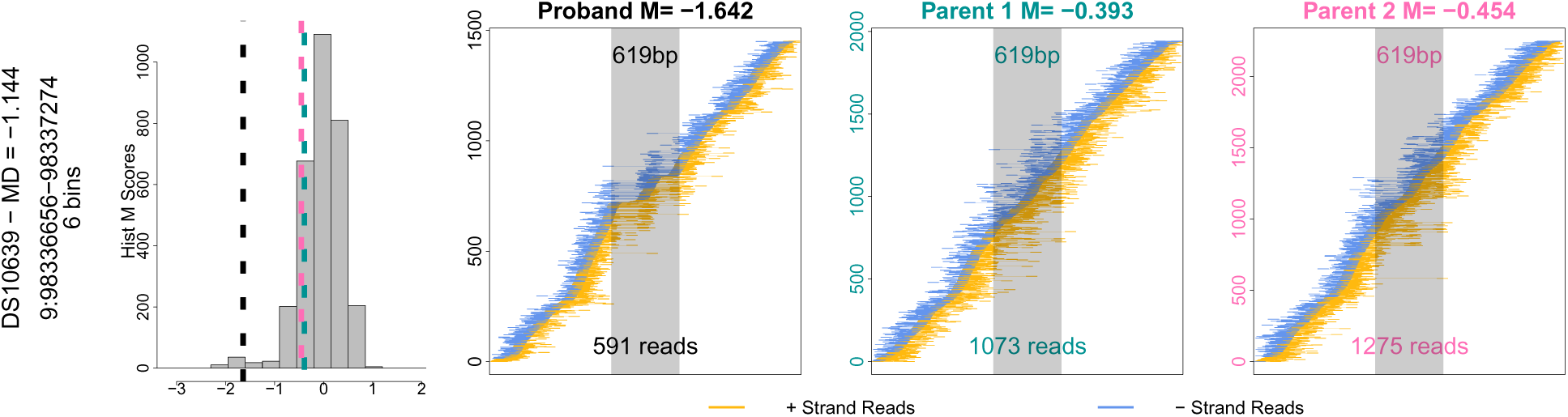
A technical (likely read mapping) artifact, resulting in a *de novo* call by CANOES:b. This pattern is observed in many samples, and reflected in the variability of the M score distribution. Although the Minimum Distance is −1.14, this region is discarded by MDTS due to the spread of the M scores.

**Supplementary Figure 7:**
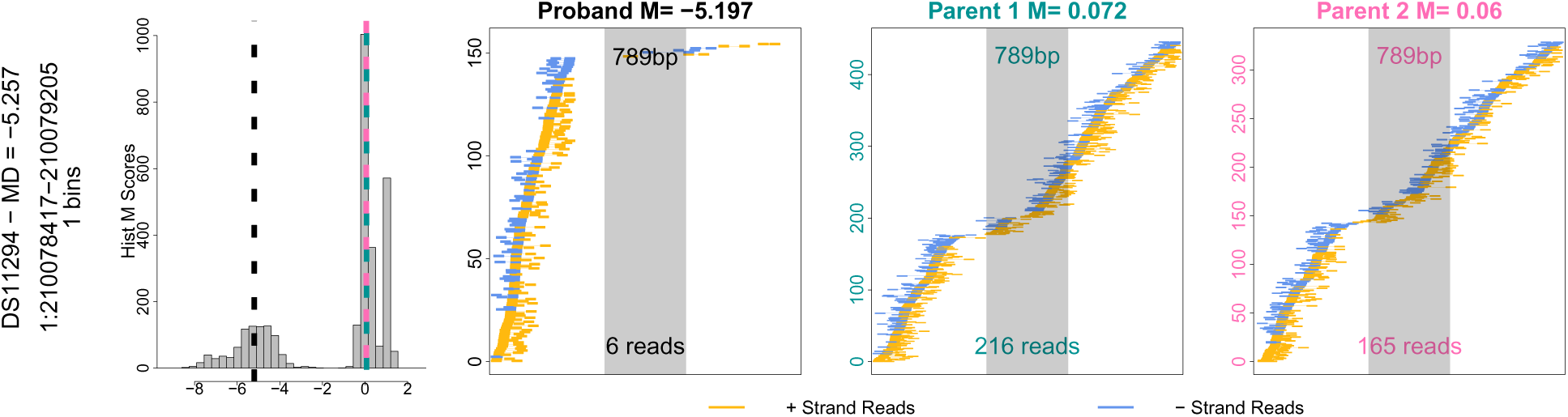
A Medelian event incorrectly called *de novo* by TrioCNV:b. The data clearly indicate a homozygous deletion in the proband, resulting through inheritance of one hemizygous deletion from each parent. This region is actually a piece of the larger CNP on chromosome 1 identified by MDTS.

**Supplementary Figure 8:**
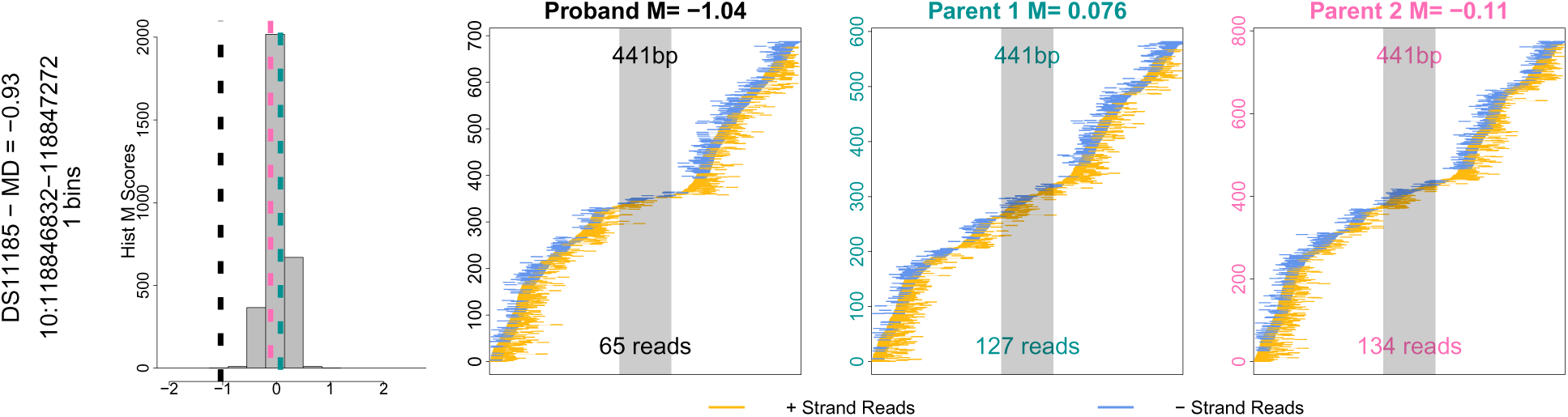
The M scores (-1.04, 0.08, and −0.11 for the proband and parents respectively) and the resulting Minimum Distance of −0.93 are consistent with a *de novo* deletion, very few reads are observed in this reqion, resulting in one bin only and highly variable statistics. In addition, no discordant read-pairs span this 441bp region.

